# Endospore Appendages Enhance *Bacillus cereus* Spore Adhesion to Industrial Surfaces, Modulated by Physicochemical Factors

**DOI:** 10.1101/2025.05.06.652425

**Authors:** Unni Lise Albertsdottir Jonsmoen, Jennie Ann Allred, Dmitry Malyshev, Jonas Segervald, Magnus Andersson, Marina Elisabeth Aspholm

**Affiliations:** Department of Paraclinical Sciences, Faculty of Veterinary Medicine, Norwegian University of Life Sciences (NMBU), Ås, Norway; Department of Physics, Umeå University, Umeå, Sweden; Umeå Centre for Microbial Research (UCMR)

## Abstract

Spores of species belonging to the *Bacillus cereus* group (*B. cereus* spp.) are common contaminants in food processing environments due to their ability to adhere to surfaces and resist cleaning procedures. These spores are equipped with pilus-like endospore appendages (ENAs), which are believed to promote surface adhesion. We have investigated the role of ENAs in spore adhesion to abiotic surfaces using wild-type (WT) *B. cereus* spp. strains and isogenic mutants lacking ENAs or an intact exosporium. WT spores expressing both short and long ENAs (S+L+) adhered significantly more to stainless steel (SS) and polypropylene (PP) compared to bald spores (S–L–) and spores of an exosporium-deficient mutant (Δ*exsY*), while adhesion to polystyrene (PS) and glass was not significantly affected by the presence of ENAs. The Δ*exsY* mutant showed the lowest adhesion across all tested surfaces, a pattern also observed for vegetative cells. The individual roles of S-ENA and L-ENA were also assessed, where the strongest adhesion occurred when both fibers were present on PP. A trend also emerged on PP; WT remained adhered for an hour, while bald spores tended to detach within that time. For saline conditions and pH, bald spores adhered strongly to SS, while for non-ionic surfactant and concentrated protein solutions, WT spores exhibited the strongest adhesion. These results highlight the crucial role of ENAs in *B. cereus* spp. spore adhesion to industrial relevant surfaces, providing mechanistic insight into spore persistence. This understanding supports the design of surface treatments to prevent contamination, spoilage, and foodborne illness.

**Importance:** Bacteria belonging to the *Bacillus cereus* group represent a persistent challenge in food production due to their highly resilient endospores (spores), which resist cleaning and disinfection as well as food processing. Understanding the adhesion properties of these spores is essential for designing effective surface treatments that reduce chemical use, enhance food safety and quality, and minimize environmental impact.

This study underscores the important role of endospore appendages (ENAs) in spore adhesion to common materials in food processing and laboratory environments. Wild-type spores expressing both S-ENA and L-ENA adhered significantly more than mutants lacking ENAs or the exosporium, highlighting ENAs as potential targets for disrupting spore adhesion. Time-dependent adhesion tests on polypropylene revealed strong, sustained attachment by wild-type spores, contrasting with weaker, transient adhesion by ENA-depleted mutants.

These findings offer valuable insights into *B. cereus* spore adhesion dynamics, guiding the development of tailored cleaning protocols to improve contamination control and sustainability.

## Introduction

Spore-forming bacteria belonging to the *B. cereus* group pose significant challenges to the food industry, frequently contaminating dairy products and other food items such as rice, pasta, spices, and vegetables (Stenfors Arnesen et al., 2008). This group of bacteria produces a range of different degradative enzymes during growth in food, with many strains also producing toxins associated with food-borne illnesses (Dietrich et al., 2021; Granum & Lund, 2006). The hygienic risk associated with these strains is amplified by their ability to produce highly resilient endospores, commonly referred to as “spores”. Spores can survive high heat and pressure, including pasteurization, desiccation, and other food processing steps that are intended to eliminate contaminating microorganisms (Wells-Bennik et al., 2016). Moreover, they can form biofilms that are particularly challenging to eradicate once established. The protective extracellular matrix of the biofilm shields embedded cells and spores from cleaning agents and environmental stresses while also facilitating periodic release of cells into the surroundings. This results in persistent contamination in food production environments, causing an ongoing threat to both product safety and quality (Seale et al., 2008). The presence of biofilm not only increases the risk of foodborne illnesses but also complicates cleaning and sanitization processes, leading to higher operational costs and waste of food due to suboptimal quality. Effective mitigation strategies require a comprehensive understanding of the mechanisms driving spore adhesion and biofilm formation.

In the food industry, food contact surfaces are often composed of stainless steel and plastic. Additionally, typical food and drink containers, as well as sample vials and disposable laboratory equipment used for contamination assessments, are often made of plastic and glass. Several studies have shown that *B. cereus* spp. spores bind effectively to these materials, with some attributing their strong adhesion to the hydrophobic nature of the exosporium (Andersson & Rönner, 1998; Rönner et al., 1990). Moreover, surface appendages and low zeta potential are believed to significantly enhance the spores’ adhesiveness (Husmark & Rönner, 1992). However, there are conflicting findings in the literature. Other studies have not established the same link to spore surface hydrophobicity, instead attributing the strong adhesiveness of *B. cereus* spores to their surface appendages (Mercier-Bonin et al., 2011; Tauveron et al., 2006). Earlier functional studies on *B. cereus* spore appendages related to their role in adhesion to abiotic surfaces were primarily done by comparing spores with intact appendages to those from which these fibers had been mechanically removed (Husmark & Rönner, 1992; Klavenes et al., 2002; Stalheim & Granum, 2001). However, these studies produced variable results, with some reports indicating enhanced adhesion, and others finding no significant correlation or even suggested that the appendages might reduce adhesion under certain conditions (Faille et al., 2010; Husmark & Rönner, 1992; Mercier-Bonin et al., 2011; Tauveron et al., 2006; Klavenes et al., 2002). These inconsistencies highlight a knowledge gap in our understanding of the mechanisms underlying spore adhesion and the functional role of their appendages.

The structure and composition of endospore appendages (ENAs) remained unresolved for a long time, but technological advancements have enabled further research to unravel their characteristics and functions. Recent studies on the foodborne outbreak strain *Bacillus paranthracis* NVH 0075/95 (classified as *B. cereus* until 2022) have identified two distinct types of ENAs: the staggered S-ENA and ladder-like L-ENA (Pradhan et al., 2021). The S-ENA protrudes from the endospore coat and is 8-12 nm thick, 0.4-1.2 µm long and displays four to five filamentous fibrillae at its distal end (Plomp et al., 2005; Pradhan et al., 2021). This fiber is built of two major protein subunits, Ena1A and Ena1B, which self-assemble into a helical structure, as illustrated in Figure 1. The L-ENA, which is thinner and shorter than the S-ENA, is attached to the exosporium layer, measures 2.5–3.5 nm in thickness, 0.2–1.6 µm in length and is built of the major subunit Ena3A (Plomp et al., 2005; Sleutel et al., 2024). Unlike the helical structure of the S-ENA, characterized by its spiral arrangement of protein subunits, the L-ENA’s morphology resembles a series of stacked disks. The L-ENA fiber also carries a single tip fibrillum at its distal end, which is composed of the collagen-like L-BclA protein, a member of the *Bacillus* collagen-like protein family (Sleutel et al., 2024). Recent findings by Jonsmoen *et al*. 2024 suggest that L-BclA is also expressed at the end of the S-ENA fiber, forming one or more of its terminal fibrillae (Jonsmoen et al., 2024).

**Figure 1:**
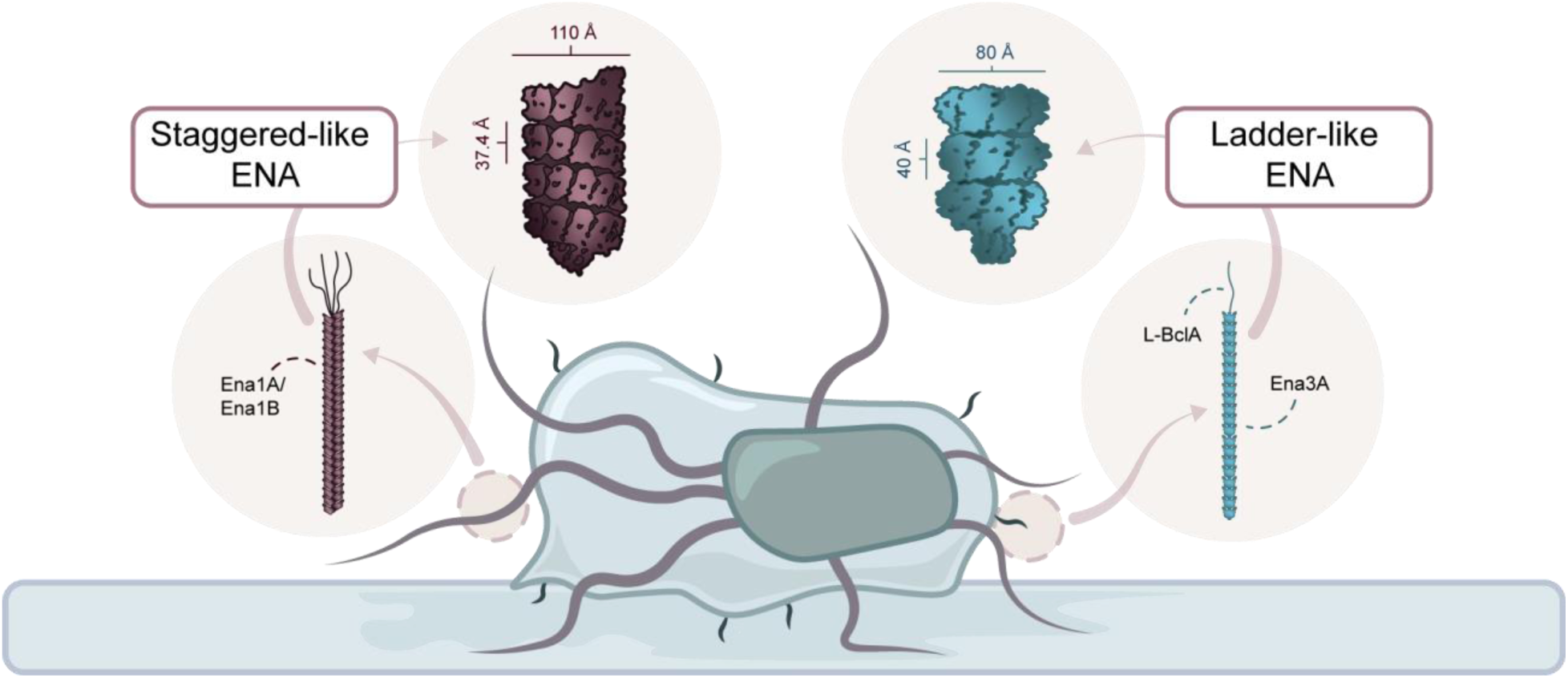
Schematic illustration of an endospore adhered to a surface, with both S-ENAs (red) and L-ENAs (blue) highlighted.

The recent discoveries about the structure and genetic origins of ENAs have enabled the development of mutants lacking S- and/or L-ENAs, allowing for more detailed studies of their function(s) (Jonsmoen et al., 2023, 2024). For example, both S- and L-ENAs have been shown to play key roles in spore-to-spore contact, leading to spore self-aggregation (Jonsmoen et al., 2024), and that appendages, particularly L-ENAs, were found to enhance adhesion to glass surfaces (Jonsmoen et al., 2023). However, the role of ENAs in binding to common materials used in food production and laboratory settings, as well as the influence of chemical conditions on their adhesive properties, remains poorly understood.

In this study, we hypothesize that S- and L-type ENAs play a role in enhancing spore adhesion to abiotic surfaces commonly encountered in food production and laboratory environments. Using *B. paranthracis* NVH 0075/95 and isogenic mutants with distinct ENA phenotypes, we evaluated spore adhesion to stainless steel (SS), commonly used in food processing equipment; polystyrene (PS) and polypropylene (PP), prevalent in packaging and sampling materials (Guazzotti et al., 2022); and glass, a material of central importance in laboratory environments. Additionally, we also examined how the presence of salt (buffer solution), pH and extended contact time influence spore adhesion.

Understanding the interaction between *B. cereus* spores, particularly the role of ENAs, and various contact materials can improve sample accuracy and lead to more effective strategies for preventing spore adhesion. This knowledge may not only improve the reliability of laboratory analyses but also support the development of more efficient approaches to control biofilm formation, thereby ensuring better food safety practices and improved shelf life of food products.

## Results

### ENAs influence spore surface adhesion to stainless steel and polypropylene

The role of ENAs in adhesion to abiotic surfaces — SS, PP, PS and glass— was assessed by comparing wild-type (WT) *B. paranthracis* NVH 0075/95 spores (S+L+) with bald spores lacking both short and long ENAs (S–L–), as well as with spores lacking a complete exosporium due to deletion of the major exosporium protein gene, *exsY*, although some residual exosporium fragments remain. The Δ*exsY* mutant, while producing an incomplete exosporium, still generates S-ENA; however, L-ENAs are absent, as they are anchored to the exosporium surface (Jonsmoen et al., 2023, 2024). To provide a broader comparison, adhesion of vegetative *B. paranthracis* cells was also measured. The vegetative cells carry flagella-like extensions (Jonsmoen et al., 2024) but do not express ENAs (Pradhan et al., 2021; Sleutel et al., 2024). The morphology and genotypes of all strains used in this study are summarized in Table 1.

**Table 1:** The genotype and morphology of the strains used in this work.

| Name | Genotype | Spore surface-related phenotype |  |  | Reference |
| --- | --- | --- | --- | --- | --- |
|  |  | S-ENA | L-ENA | Exosporium |  |
| S+L+ (WT) | WT | + | + | + | (Lund & Granum, 1996) |
| S+L- | *Δena3* | + | - | + | (Jonsmoen et al., 2023; Sleutel et al., 2024) |
| S-L+ | *Δena1ABC* | - | + | + | (Pradhan et al., 2021) |
| S-L- (bald) | *Δena1ABC*<br>*Δena3* | - | - | + | (Jonsmoen et al., 2023; Sleutel et al., 2024) |
| *ΔexsY* | *ΔexsY* | + | - | - | (Jonsmoen et al., 2023) |

The role of ENAs on spore adhesion to SS, PP, PS and glass was evaluated by sequentially transferring spores and vegetative cells between vials or tubes and then measuring the reduction in OD600, which served as an indicator of spore loss due to adherence to the materials tested (Figure 2A). The results demonstrated that ENA-mediated spore adhesion was surface-dependent (Figure 2B). On SS, wild-type spores exhibited the highest adhesion efficiency, while both bald and Δ*exsY* spores showed significantly reduced adhesion compared to those of the WT strain (*p* = 0.0014 and *p* = 0.0003, respectively, after 10 transfers). A similar pattern was observed on PP, where WT spores adhered significantly more than both bald spores (*p* = 0.0004) and exosporium-deficient spores (*p* = 0.0002). In contrast, adhesion to PS and glass was equally efficient for WT and bald spores, suggesting that ENAs play a less prominent role in spore attachment to these surfaces. Nonetheless, Δ*exsY* spores again displayed the lowest adhesion on both glass (*p* = 0.0005) and PS (*p* = 0.0002) after 10 transfers, suggesting that an intact exosporium may contribute to adhesion independently of ENAs on certain surfaces.

**Figure 2:**
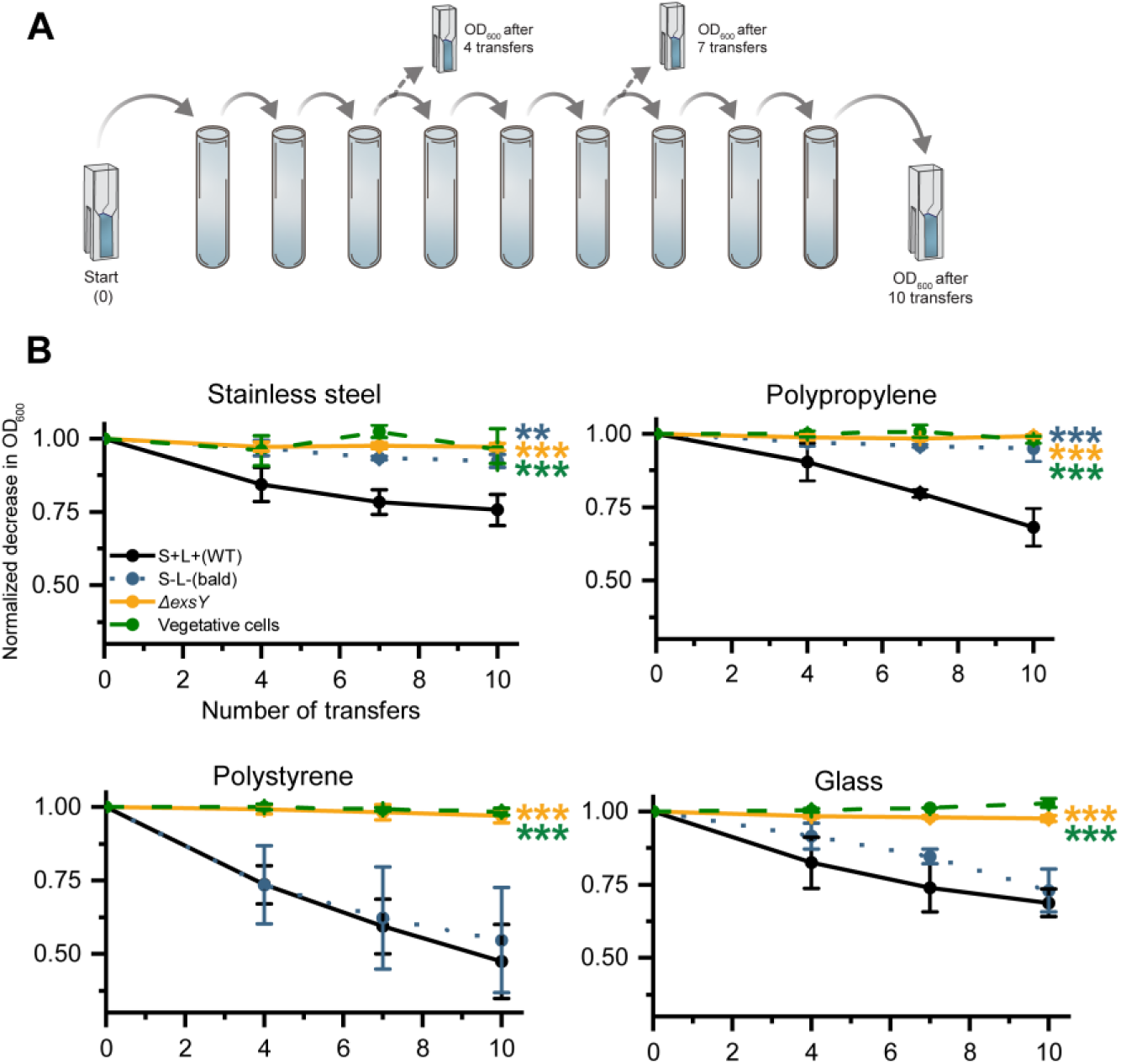
**(A)** Schematic illustration of the methodology for measuring spore and cell adhesion to stainless steel, polypropylene, polystyrene and glass by loss of OD_600_ after 4 transfers, 7 transfers and 10 transfers. **(B)** The measured adhesion of WT, bald, Δ*exsY* spores and vegetative cells, on SS, PP, PS and glass, revealing varied impact of ENAs across the different tested materials. The largest impact of ENAs was attributed to SS and PP. Exosporium-deficient spores and vegetative cells had the least adhesion efficiency for all materials tested. One asterisk (p ≤ 0.05) indicates significant difference, with two (*p* ≤ 0.01) and three (*p* ≤ 0.001) asterisks denoting increasingly lower p-values.

Adhesion of vegetative cells to the same materials were also assessed. These cells showed minimal adhesion across all tested surfaces, with levels even lower than those of the *ΔexsY* spores (Figure 2B). This underscores the limited adhesive capacity of vegetative cells and highlights the distinct contribution of the exosporium and ENAs to spore adhesion in *B. paranthracis*.

Overall, a clear trend emerged: WT spores adhered more efficiently than bald spores, Δ*exs*Y spores, and vegetative cells on SS and PP. On PS and glass, however, WT spores only outperformed Δ*exsY* spores and vegetative cells, with no significant difference for bald spores. These findings underscore the complex, surface-specific role of ENAs in spore adhesion, suggesting that their contribution is strongly influenced by the physicochemical properties of the substrate.

### Limited role of surface and spore hydrophobicity in spore adhesion

Since ENAs significantly influenced spore adhesion to both SS and PP, as demonstrated by comparisons between wild-type spores and those lacking both S- and L-ENAs, we next investigated whether hydrophobicity could help explain these patterns. To assess potential differences in hydrophobicity between the materials we performed sessile drop contact angle measurements using water as the test liquid. By measuring the angle formed between the liquid droplet and the surface material, we could determine if the surfaces were hydrophobic or hydrophilic. As seen in Figure 3, both PP and PS exhibited contact angles greater than 90°, classifying these as hydrophobic. In contrast, SS showed a contact angle below 90°, indicating a hydrophilic surface. Accurate measurement of wettability on glass was not possible due to the curvature of the substrate, only its outline is included in the figure. Raw data are available in Supplementary Information under data for contact angle and surface roughness measurements, Figure S1 and Table S1.

**Figure 3:**
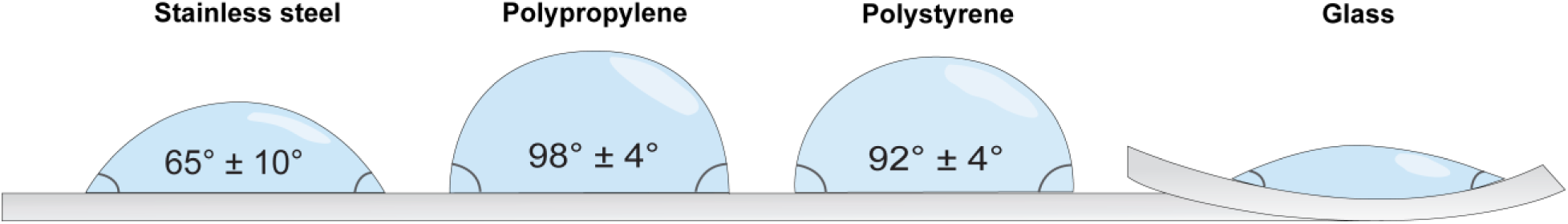
Contact angles were determined by measuring the angle formed between a water droplet and the different surface materials (SS, PP, PS and glass). The resulting droplet profiles are presented as an outline for all materials. The contact angle for glass could not be determined due to the highly curved surface, which resulted in unreliable measurements. SS exhibited hydrophobic properties, while PP and PS were classified as hydrophilic.

Because of the varying wettability of the materials, isolating the specific effect of hydrophobicity on spore surface adhesion was not possible. Therefore, to quantify adhesion under controlled conditions, we adapted the spore transfer setup to 96-well microtiter plates with uniform well dimensions and measured absorbance at 600 nm. This setup allowed us to compare spore adhesion on hydrophilic surfaces (PS-treated) with a hydrophobic surface (PS-untreated) (Spriano et al., 2017; Thermo Scientific, n.d.). The two types of PS materials were selected to provide a controlled baseline for surface hydrophobicity, while minimizing the potential influence of other factors, e.g., surface roughness and hardness.

The results presented in Figure 4 show that WT spores do not preferentially adhere to either the hydrophobic or hydrophilic surface, as no statistically significant difference was observed after the tenth transfer (*p* > 0.1). Furthermore, the presence of ENAs did not appear to influence adhesion, as both WT and bald spores adhered with similar efficiency to both surfaces. However, bald spores displayed significantly higher adhesion to the hydrophilic surface compared to the hydrophobic substrate after ten transfers (*p* = 0.0470). Consequently, the inherent wettability of the substrate does not appear to be the primary factor dictating ENA-mediated spore adhesion. Instead, the spores’ intrinsic properties—potentially mediated by ENAs—seem to play a more decisive role.

**Figure 4:**
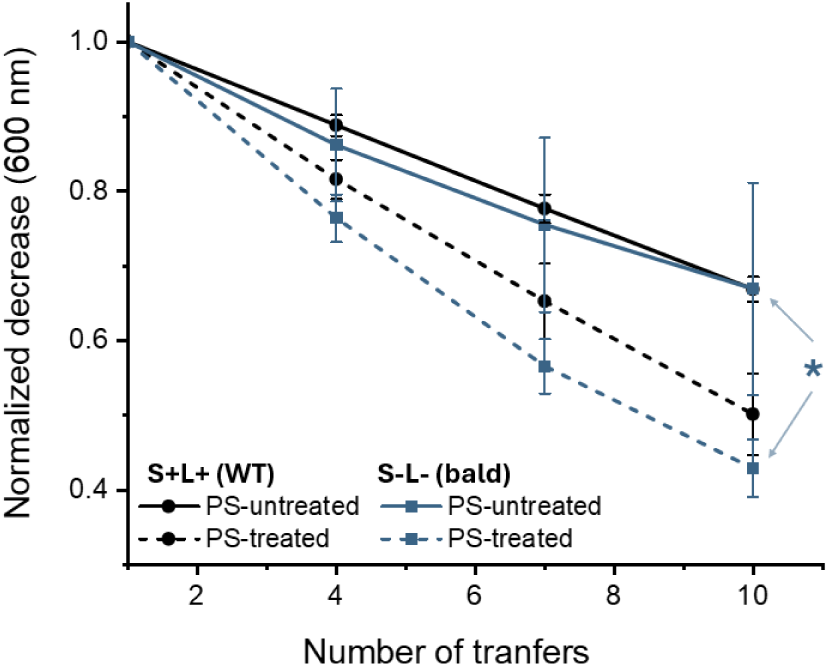
Adhesion efficiency of WT and bald spores after 10 transfers was assessed on polystyrene (PS) surfaces with different treatments: untreated (hydrophobic) and treated (hydrophilic). The results showed that WT (S+L+) spores exhibited no significant preference between the two surface types. In contrast, bald (S–L–) spores adhered significantly more to the hydrophilic surface, as indicated by a single asterisk denoting statistical significance (*p* ≤ 0.05). A significant difference was also observed between WT spores on the treated surface and bald spores on the untreated surface; however, this comparison is not shown in the figure, as it falls outside the primary scope of this study.

We also investigated whether ENAs affect spore surface hydrophobicity by performing a microbial adhesion to hydrocarbon (MATH) test. The results showed no significant differences in hydrophobicity between ENA-depleted spores and WT spores (Figure 5), although S-L- spores displayed a relatively high standard deviation of 16%. In contrast, the exosporium-deficient mutant exhibited significantly lower hydrophobicity compared to both WT and ENA-depleted spores (*p* < 0.0001), indicating that the exosporium structure, rather than ENAs, play a key role in modulating spore surface hydrophobicity.

**Figure 5:**
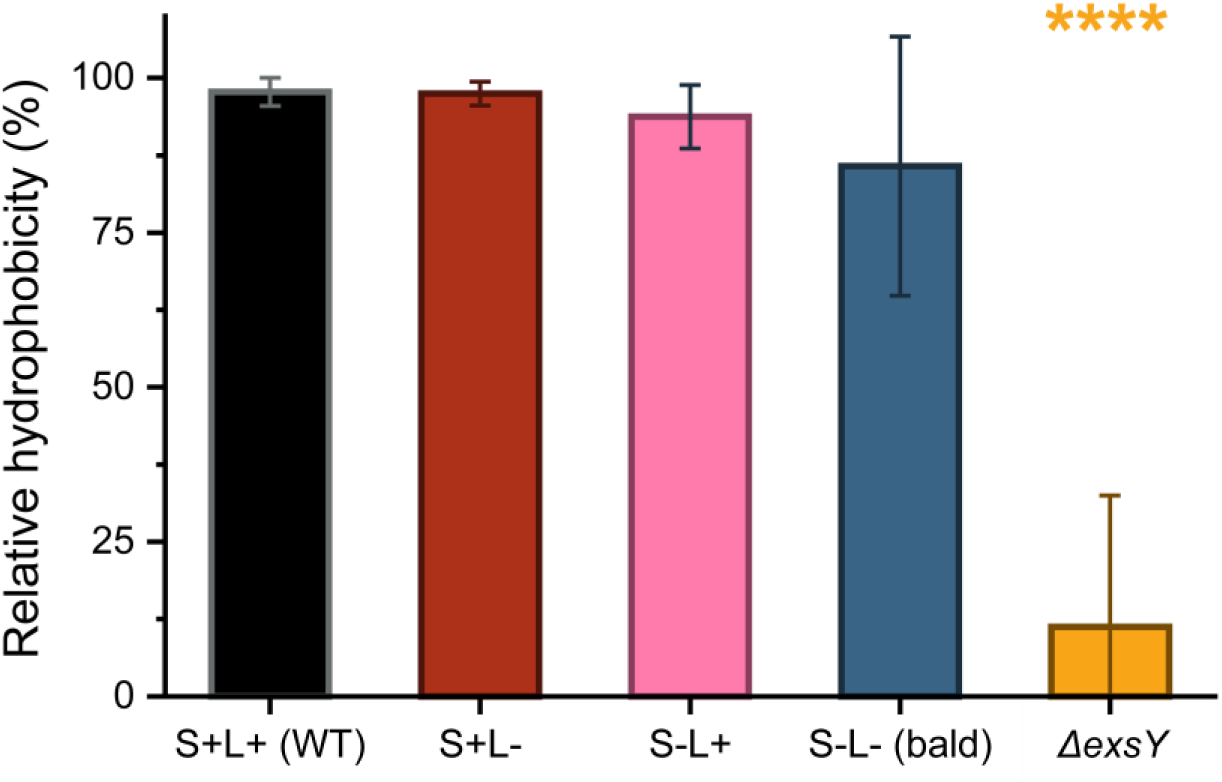
Relative hydrophobicity of WT spores was compared to various isogenic mutant spores, using the MATH test. The results show that removal of the ENAs does not significantly affect the hydrophobicity of the spores. However, the exosporium-deficient spores exhibited the lowest relative hydrophobicity compared to WT, with a statistical significance *p* < 0.0001 as denoted by the four asterisks in the figure. The results are based on 3 biological triplicates.

In conclusion, neither surface nor spore hydrophobicity appear to influence adhesion to the various substrates mediated by ENAs. To better understand why ENAs promote strong adhesion to some materials but have a lesser effect on others, we further investigated the intrinsic properties of the materials.

### Stainless steel features large surface structures, while glass exhibited overall lowest roughness

To better understand how the inherent properties of the tested substrate influence spore adhesion, a detailed material characterization was performed using scanning electron microscopy (SEM), tensiometer, profilometer, and atomic force microscope (AFM). This in-depth analysis provides insights into the factors driving preferential spore adhesion to specific substrates.

As shown in Figure 6A, the SEM micrographs revealed that the surfaces PP, PS, and glass were relatively smooth and uniform, with minimal irregularities. In contrast, the SS exhibited a highly textured surface characterized by grooves and ridges. To quantify these differences in surface topography, AFM and profilometry were performed. Surface scanning with a sharp probe revealed that glass had the lowest roughness (5 nm ± 2 nm), followed by PS (29 nm ± 16 nm), PP (92 nm ± 20 nm), and SS (7 µm ± 0.4 µm), which displayed the highest roughness as determined by root mean square (RMS) roughness measurements. Surface topography was also visualized as height maps in Figure 6B, where darker regions represent lower areas and brighter regions indicate elevated areas. These visualizations further supported the calculated RMS roughness, confirming that SS had the roughest surface while glass had the smoothest. Additionally, the surface ratio (SR) was calculated to assess how surface texture contributes to an increase in actual surface area relative to the scanned area. Both glass and PS exhibited surface ratio values close to 1, indicating minimal surface expansion due to texture. In contrast, SS and PP showed slightly elevated surface ratio values (> 1), suggesting a larger increase in actual surface area because of surface roughness. Lastly, autocorrelation analysis was performed to further understand the repetitive distribution of the grooves and ridges on each material surface. A Fourier transform was used to identify dominant periodic features independent of direction, while both vertical and horizontal autocorrelations provided insight into overall surface patterns, see tables S2-5 in Supplementary Information under data contact angle and roughness measurements for details. Overall, SS exhibited larger-scale periodic features, as indicated by both Fourier and spatial autocorrelations analyses, whereas PP, PS and glass displayed finer more closely spaced surface patterns. Interestingly, as seen on the height map, polypropylene also exhibited a pronounced directional roughness.

**Figure 6:**
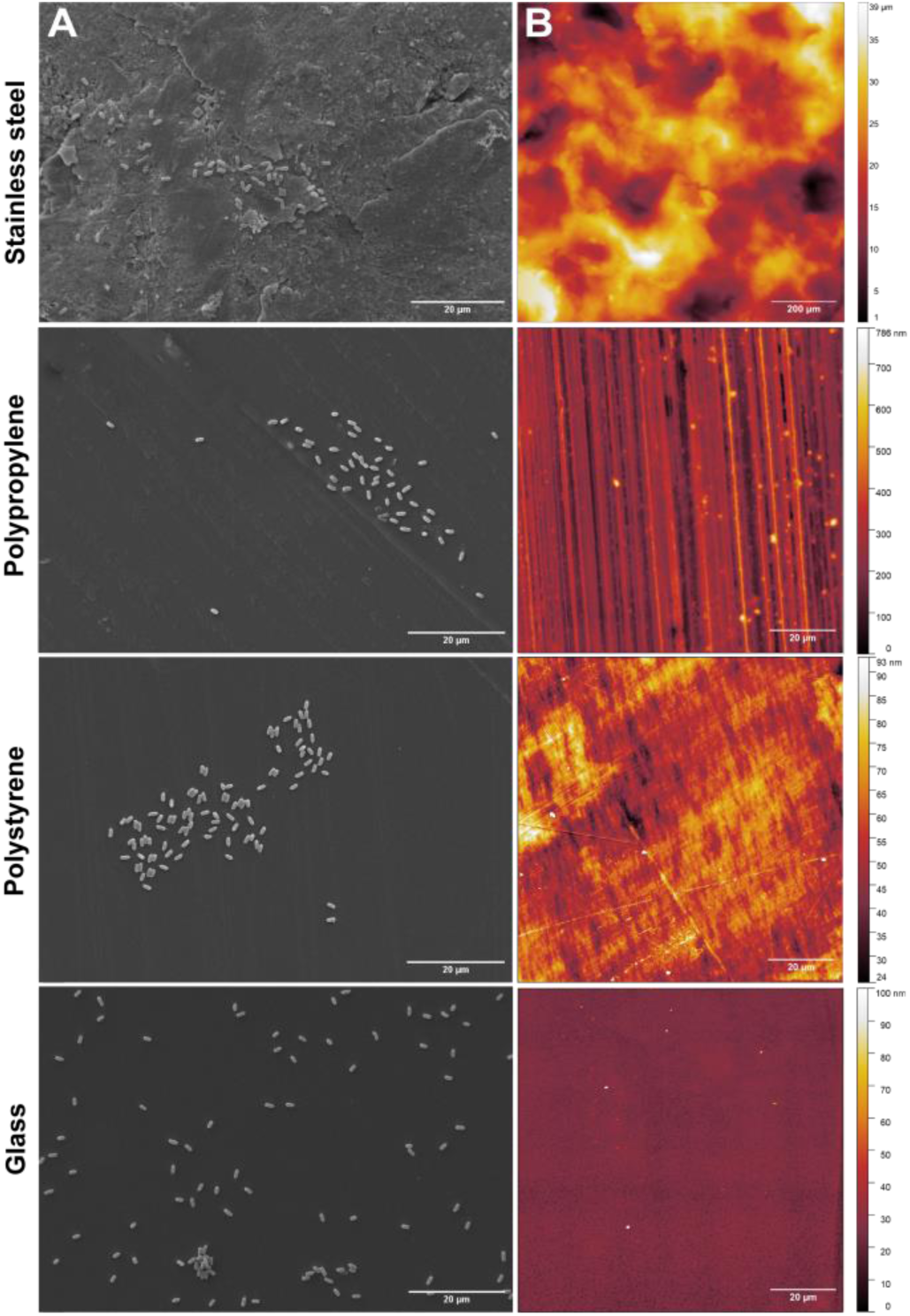
**(A)** Scanning electron micrographs showing spores adhered to stainless steel, polypropylene, polystyrene and glass surfaces. The SS surface showed pronounced irregularities, in contrast to the glass surface, which appeared smoothest overall. A scale bar of 20 µm is shown for all materials. **(B)** Height maps were generated using a profilometer for stainless steel and AFM for PP, PS and glass. The range of surface heights is displayed on the right side of each map, presented in µm for SS and in nm for PP, PS, and glass.

### Synergistic effect of S-ENAs and L-ENAs essential for spore adhesion to polypropylene

Recognizing the importance of ENAs in spore adhesion, two additional isogenic mutant strains were included to elucidate the distinct roles of S-ENAs and L-ENAs. These strains lack either S- ENA (S-L+) or L-ENAs (S+L-), enabling investigation of the contribution of each appendage type to the adhesion process. WT spores, bald spores, exosporium-deficient spores, and vegetative cells were included in the analysis for comparison. PP and SS were selected as substrates, as previous experiments (Figure 1B) showed a significant difference in adhesiveness between WT and bald spores on these materials. Consistent with earlier results, a significant difference in adhesion to PP was observed between WT spores, bald spores, exosporium-deficient spores, and vegetative cells, with WT showing the highest adhesion after 10 transfers (Figure 7). Additionally, both S+L- and S-L+ spores showed significantly reduced adhesion compared to WT spores (p=0.0114 and p=0.0031, respectively). These findings indicate that both S-ENAs and L-ENAs are necessary for spores to adhere effectively to polypropylene surfaces. Similarly, on SS, WT spores adhered significantly more than bald, exosporium-deficient spores and vegetative cells after 10 transfers. However, in contrast to the results on PP, no significant difference was observed between WT, S+L- and S-L+ spores on SS. This suggests that, on SS, the presence of either S-ENA or L-ENA alone is sufficient to achieve an adhesion efficiency comparable to WT spores. These findings highlight the collaborative role of both S-ENAs and L-ENAs, suggesting that both appendage types are essential and act collectively to mediate effective spore attachment to certain abiotic surfaces. However, in the case of SS, the presence of either S-ENA or L-ENA alone was sufficient to achieve optimal adhesion.

**Figure 7:**
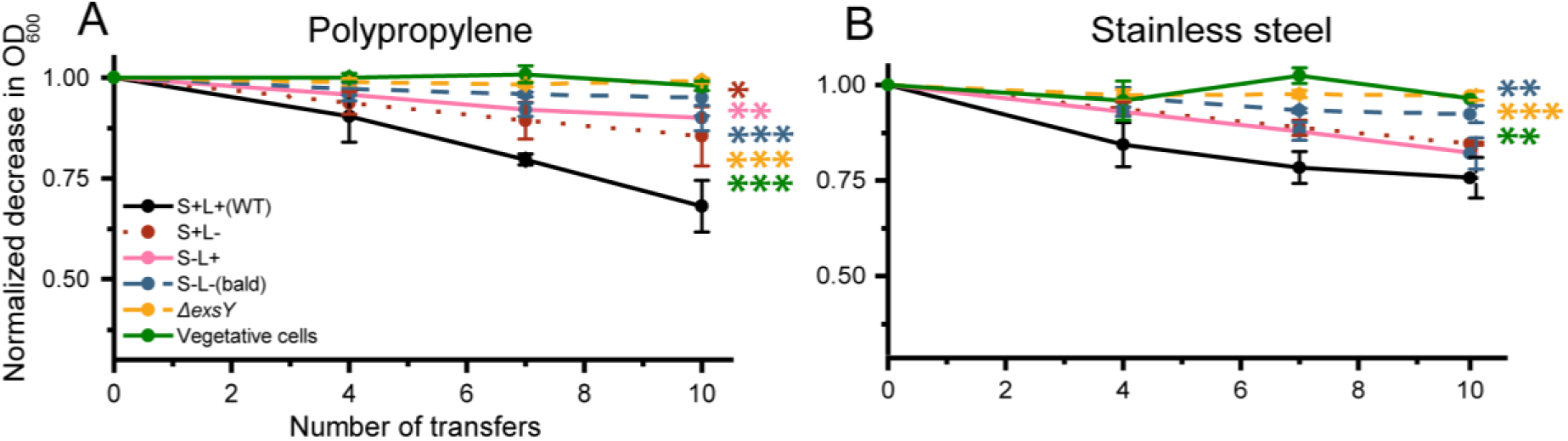
Loss of OD_600_ during sequential transfer of spores, with and without appendages, as well as isogenic mutant strains expressing only S-ENA (S+L-) or only L-ENA (S-L+), measured between PP tubes (A) and SS coupons (B). **(A)** After 10 transfers on PP, spores lacking either S-ENAs or L-ENAs showed significant reduction in adhesion compared to WT spores, indicating a collaborative role of both appendage types in achieving optimal adhesion. **(B)** In contrast, no significant difference was observed between WT and either mutant strain on SS, suggesting that the presence of either S-ENA or L-ENA alone is sufficient for effective adhesion to this surface. One asterisk (*p* ≤ 0.05) indicates significant difference, with two (*p* ≤ 0.01) and three (*p* ≤ 0.001) asterisks denoting increasingly lower p-values.

### ENAs may be crucial for maintaining adhesion to polypropylene over time

Previous experiments demonstrated substrate-dependent variations in spore adhesion, with ENAs playing a critical role, particularly on PP and SS. These measurements were taken over relatively short time periods, typically only a few seconds per transfer step. In industrial environments, spores are exposed to both continuous and intermittent flows (“stop and go”), resulting in varying contact times on the abiotic surfaces. To understand the effect exposure time has on spore adhesion, we conducted an extended incubation experiment using PP tubes, allowing us to also implement rotation to emulate fluid conditions as observed in industry. WT and mutant spores were incubated and continuously rotated for 1, 10, 30, and 60 minutes. Adhesion was quantified by measuring the reduction in OD600 of the spore suspension after each time point (Figure 7).

The results revealed a clear pattern of adhesion dynamics over time. WT spores and with spores expressing either S-ENA or L-ENAs maintained the highest level of adhesion throughout the 60-minute incubation, demonstrating strong and sustained attachment to the PP surface. In contrast, the exosporium-deficient mutant exhibited consistently weak adhesion at all points, underscoring the importance of the exosporium and its associated appendages in mediating stable surface interactions. Bald spores, lacking both S- and L-type ENAs, displayed a more transient adhesion profile. Their adhesion peaked at 30 minutes, suggesting an initial ability to bind effectively to the PP surface. However, by the 60-minute mark, their adhesion level decreased sharply, aligning with those of the exosporium-deficient mutant. This significant drop in adhesion indicates that bald spores rely on transient binding mechanisms, which are insufficient for maintaining long-term attachment to PP surfaces. Although statistical significance could not be confirmed at either the 30- or 60-minute mark due to due to large standard errors, the observed trend supports the conclusion that ENAs play a critical role in sustaining spore adhesion over extended periods, particularly on substrates like PP.

### Ionic concentration impacts spore adhesion to stainless steel

Previous work demonstrated that the presence of salt in a spore suspension inhibits appendage-mediated spore-spore interactions (Jonsmoen et al., 2024). To investigate if similar effects occur in spore adhesion to abiotic surfaces, we examined the influence of salt and other relevant factors in the liquid environment. Spores were suspended in phosphate-buffered saline (PBS) to simulate elevated ionic strength; in NaOH3 and HNO3 to model chemical conditions during disinfection and cleaning-in-place procedures; in Tween-20 to represent high concentrations of non-ionic surfactants; and in bovine serum albumin (BSA) to mimic environments with increased protein content. SS was used as the sole surface due to its relevance in industrial settings.

WT spores exhibited consistent adhesion to SS surfaces regardless of pH, showing no significant difference between acidic (0.1 M HNO₃, pH 1.7) and alkaline (0.1 M NaOH, pH 13.8) conditions compared to adhesion in sterile water. In contrast, bald spores displayed significantly increased adhesion under both acidic and alkaline conditions, with *p* = 0.0001 and *p* = 0.0008, respectively, after 10 transfers (Figure 9A–B). A similar pattern was observed in phosphate-buffered saline (PBS), where bald spores adhered more effectively under elevated ionic strength (0.5× and 1× PBS) compared to sterile water, with *p* = 0.0169 and *p* = 0.0123, respectively (Figure 9C–D). Notably, increased salt concentration did not influence adhesion of WT spores, which remained constant across all tested conditions.

**Figure 8.**
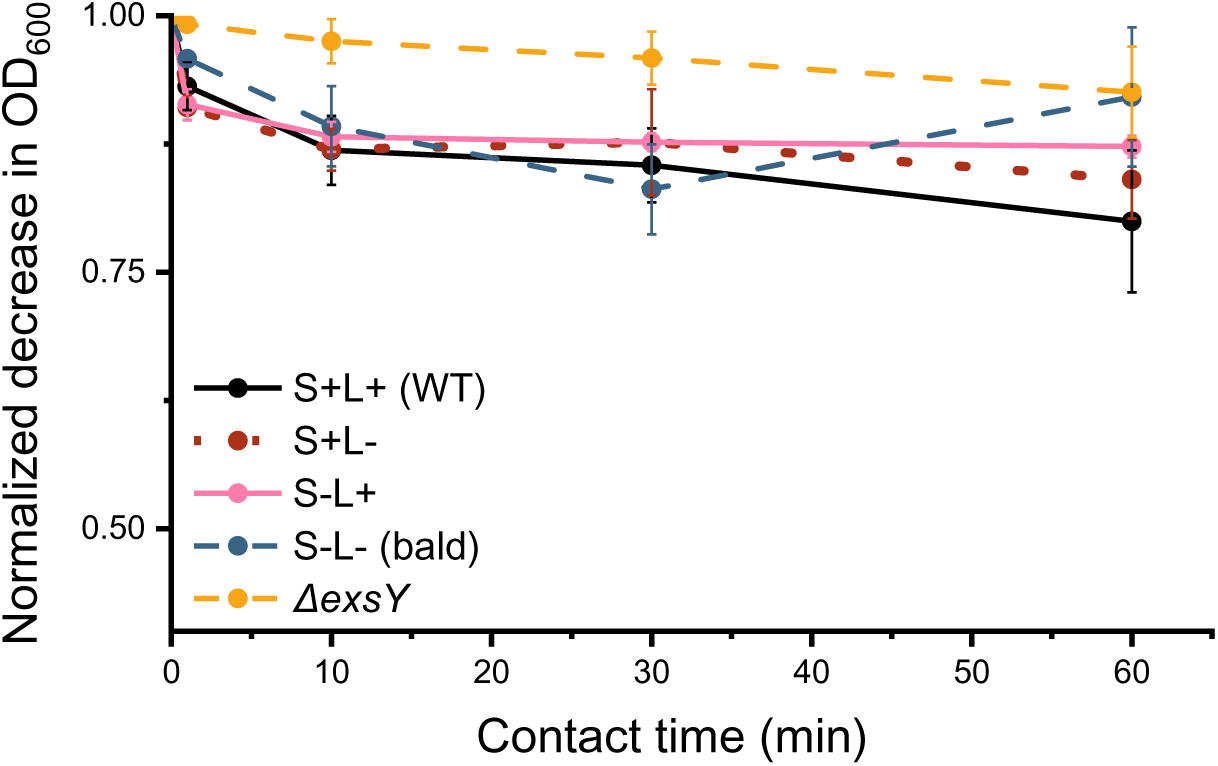
Adhesion of WT and isogenic mutant spores to PP was assessed at various incubation time points (1 minute, 10 minutes, 30 minutes and 60 minutes). The results showed that bald spores exhibited transient adhesion efficiency, with a peak at 30 minutes followed by a decline after 60 minutes. In contrast, WT spores maintained stable adhesion over time. Data represents three biological replicates.

**Figure 9.**
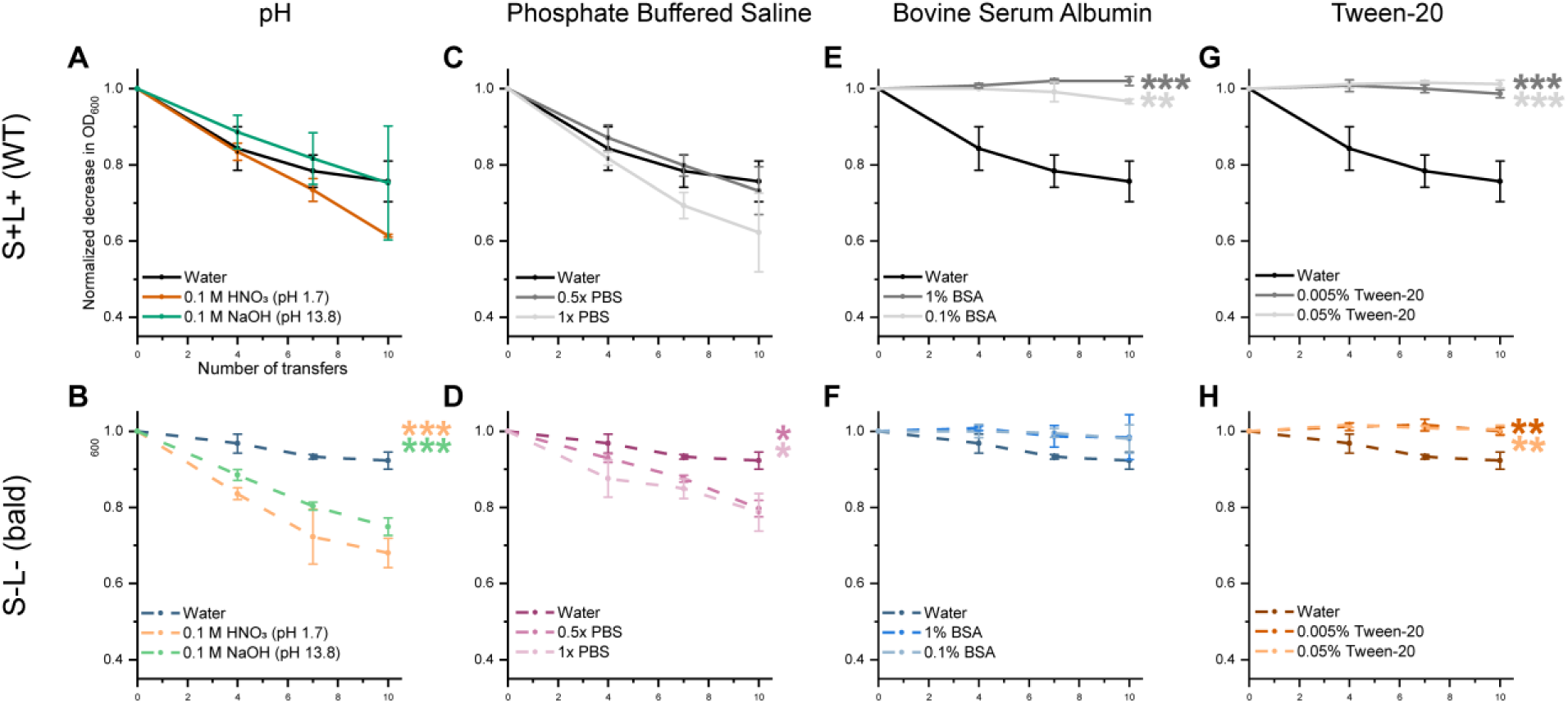
Loss of OD_600_ during sequential transfers between SS coupons is shown for WT spores (solid lines) and bald (dashed lines) spores suspended in sterile water and under different other conditions: **(A-B)** 0.1 M HNO3 (pH 1.7) and 0.1 M NaOH (pH 13.8), **(C-D)** 0.5x and 1x PBS, **(E-F)** 0.1% and 1% BSA and lastly **(G-H)** 0.005% and 0.05% Tween-20. The addition of ions—either through pH adjustment or increased salt concentration (PBS)—enhanced adhesion of bald spores to the SS surface. In contrast, both BSA and Tween-20 significantly reduced adhesion for both spore types. One asterisk (*p* ≤ 0.05) indicates significant difference, with two (*p* ≤ 0.01) and three (*p* ≤ 0.001) asterisks denoting increasingly lower p-values.

In contrast to the trend observed when adding ions to the solution, the addition of BSA (Figure 9E-F) reduced the adhesion of WT at both tested concentrations (p-values ≤ 0.001), while having no significant effect on the adhesion of bald spores. This is likely due to BSA blocking available binding sites on the spore surface or the SS surface. The non-ionic surfactant Tween-20 also led to a significant reduction in adhesion for both WT and bald spores at concentrations of 0.005% and 0.05% (*p* ≤ 0.005), compared to spores suspended in sterile water (Figure 8G–H). Similarly, as for BSA, this effect may be attributed to Tween-20 saturating or masking binding sites on the SS and spore surface, thereby inhibiting spore attachment.

## Discussion

The mechanisms underlying bacterial and spore attachment to surfaces are inherently complex and influenced by a combination of environmental conditions, surface characteristics, and microbial structural components. In this study, we investigated the role of ENAs in spore adhesion to abiotic surfaces using a single strain of *B. paranthracis* and its isogenic mutants under various conditions. We further examined how selected environmental factors modulate spore adhesion under varying conditions. Our results showed that both WT and bald spores adhered most efficiently to PP and SS surfaces, whereas exosporium-deficient spores and vegetative cells exhibited significantly lower levels of adhesion. Moreover, spores with appendages adhered significantly more effectively to PP and SS surfaces compared to bald spores (Figure 2B). In contrast, no statistically significant difference (*p* > 0.05) in adhesion between WT and bald spores was observed on glass and PS, indicating that ENAs did not influence attachment to these materials.

The surface properties of the tested materials exhibit unique characteristics that significantly influence spore adhesion. For example, PP and PS are nonpolar and hydrophobic, whereas SS and glass are hydrophilic as seen with contact angles above and below 90°, respectively (Figure 3). As shown in Figure 6A, the tested surfaces also differ significantly in their structure. The role of surface roughness in bacterial adhesion is complex; while some studies suggest that increased roughness facilitates adhesion by providing more contact points or protective niches, other report minimal or negligible effects (Bohinc et al., 2014; Cai et al., 2019; Mu et al., 2023). Nevertheless, there is a general agreement that rougher surfaces gives a greater surface area, which may provide more sites for attachments as well as larger cavities that shield bacteria from hydrodynamic shear forces (Renner & Weibel, 2011; Song et al., 2015). As shown in Figure 6B, the SS surface is markedly rougher than the other materials tested, exhibiting surface features on a scale approximately 100-fold greater and with dominant structures significantly larger than bacterial endospores (1-2 µm). PP also had considerable surface roughness, albeit less pronounced than SS, but greater than that of glass and PS. Surface roughness measurements confirmed that both SS and PP have increased surface areas compared to PS and glass. Given that WT spores adhered more efficiently than bald spores on the two roughest materials, we hypothesize that ENAs particularly enhance spore adhesion to surfaces with rougher surface topographies.

While we can assess the role of ENAs by comparing adhesion of WT and mutant spores across different materials, we must be cautious when interpreting overall differences in spore adhesion between material types. The material properties may differ in additional ways not assessed in this study e.g., surface charge, topological patterns and stiffness. As our focus is specifically on the role of ENAs in adhesion, a comprehensive material characterization falls outside the scope of this work and was therefore not included. In addition, the substantial standard deviations observed in roughness, surface ratio and autocorrelation measurements likely reflect inherent variability introduced during the manufacturing process of the materials and should be considered when interpreting the results.

Consistent with previous studies, we found that spores with an exosporium adhered more efficiently to various abiotic surfaces than those with a deficient exosporium (Faille et al., 2007; Jindal & Anand, 2018). This enhanced adhesion is often attributed to the increased hydrophobicity of spores with an intact exosporium, which has been correlated with increased adhesion to hydrophobic surfaces (Faille et al., 2002, 2010; Rönner et al., 1990; Simmonds et al., 2003). However, several studies have questioned the strength of this correlation, suggesting that morphological factors, such as presence of surface structures like ENAs, may also significantly influence spore adhesion (Faille et al., 2010; Lequette et al., 2011; Tauveron et al., 2006). Our results support this view, as the hydrophobicity of the tested materials had limited impact on the adhesion of wild-type spores, whereas bald spores exhibited increased adhesion to hydrophilic (treated) PS (Figure 4).

We have previously shown that removal of S-ENAs significantly reduces the hydrodynamic diameter of the spores, thereby influencing their behavior and interactions in an aqueous environment. The presence of S-ENAs also appears to act as a steric hindrance, limiting the ability of spores to form close interactions with surfaces or other particles (Jonsmoen et al., 2023, 2024). Polystyrene is an aromatic polymer composed of a linear hydrocarbon backbone with a benzene ring attached to every second carbon atom. In this study, surface-treated polystyrene was used in an attempt to isolate the effect of hydrophobicity on spore adhesion. However, the surface modification process may also affect other material properties. For instance, chemical treatments designed to enhance hydrophilicity often introduce new functional side chains, which can alter the surface’s topographical features and physicochemical characteristics (Lackner et al., 2009). These alterations in surface structure, when considered together with the size difference between WT and bald spores, may help explain the observed differences in adhesion. Specifically, the larger size and stiffness of the ENAs on WT spores may hinder close interactions with hydrophilic surfaces by limiting contact or preventing the spore body from settling into the surface’s microfeatures. In contrast, bald spores are more compact and may form more direct and stable contact points with the hydrophilic substrate, resulting in stronger adhesion under the same conditions.

Tauveron et al., 2006, reported that spores with longer appendages adhere better to abiotic surfaces than those with shorter ones (Tauveron et al., 2006). Based on this, we hypothesized that spores expressing S-ENAs would adhere more effectively than those expressing only L-ENAs or bald spores. Consistent with this, bald spores showed significantly reduced adherence to PP and SS compared to WT spores (Figure 7, *p* = 0.0004 and *p* = 0.0065, respectively). However, for S-L+ and S-L+ spores, reduced adhesion reached statistical significance only for PP, not for SS. This suggests that expression of either ENA type alone may be sufficient to maintain WT-like adhesion to SS, whereas depletion of one appendage type disrupts adhesion, particularly on PP.

These findings underscore the complex interplay between surface material properties and bacterial surface structures and point to a complementary role of both S- and L-ENAs in facilitating efficient spore attachment to abiotic surfaces. We have previously shown that both S- and L-ENAs facilitate spore aggregation in two distinct ways: S-ENAs facilitate longer-range interaction whereas L-ENAs mediate more intimate close-range aggregation of spores (Jonsmoen et al., 2024). In the current study, we were unable to identify any differences in adhesion behavior between the two types of appendages.

Adhesion testing on PP over different incubation times revealed the dynamic nature of spore attachment and underscored the critical role of ENAs in maintaining stable interactions with abiotic surfaces. Although no statistically significant differences were observed between the strains at individual timepoints, the overall adhesion dynamics showed that WT spores consistently exhibited the highest and most stable adhesion to PP, emphasizing the importance of ENAs in sustaining prolonged surface binding (Figure 8). In contrast, bald spores exhibited a transient adhesion pattern, suggesting a reduced capacity to maintain long-term attachment. This likely reflects weaker initial binding forces that are insufficient to withstand extended incubation. These findings emphasize the time-dependent nature of spore adhesion and demonstrate the distinct contributions of ENAs and the exosporium to stable and sustained surface interactions.

A significant difference in the time-dependent adhesion study between bald and WT spores could not be confirmed, primarily due to the high variability in the adhesiveness of bald spores, which resulted in a large standard deviation. Furthermore, the MATH assay also revealed a significantly greater variability in the hydrophobicity of the bald spores compared to the other strains tested. This therefore suggests that there might be a greater variation in properties between batches of bald spores, than of strains expressing ENAs. These variations might be due to the ENAs ability to adsorb extracellular materials, such as environmental DNA, polysaccharides, and proteins. By depleting the spores of ENAs, we may limit the spore’s ability to hold onto these polymeric substances from the milieu, which might affect the dynamics facilitating spore adhesion, and introduce a grater variability.

Understanding the adhesion dynamics of ENAs provides crucial insights into how spores interact with surfaces, but adhesion is also influenced by various external factors in the surrounding solution. For instance, WT spores suspended in surfactants (e.g., Tween-20), or in solubilized protein (e.g., BSA) showed reduced adhesion, when compared to spores suspended in water (Figure 9 E-H). This is, as previously mentioned, attributed to the blocking of binding sites on both the spores and the materials, which limits interactions and thereby lowers adhesion as demonstrated by our observations. We also showed that the presence of PBS in the spore suspension resulted in a concentration-dependent reduction in both ENA- and exosporium-mediated spore-aggregation (Jonsmoen et al., 2024). The reduction was more pronounced for WT spores than for bald spores, indicating that appendage-mediated spore aggregation is more sensitive to increased ionic strength than aggregation primarily mediated by the exosporium (Jonsmoen et al., 2024). In the present study, we observed that the adhesion efficiency of bald spores to SS increased under conditions of elevated ionic strength (PBS, 0.1M HNO₃, and 0.1M NaOH), (Figure 9 A-D) whereas the adhesion of WT spores remained largely unaffected. These results are consistent with previous studies reporting enhanced spore adhesion to SS in high ionic strength conditions. This effect is thought to occur because, under high salt or acidic conditions, water is drawn away from the spore surface. This “dehydration” of the spore’s outer layer reduces the natural repelling forces that usually keep spores and surfaces apart in liquid. With these barriers weakened, spores can get closer to the surface, leading to stronger adhesion (Husmark & Rönner, 1990). In addition, bacterial appendages, like ENAs, can facilitate bacterial adhesion, as they are able to reach through the repulsive forces of the electrostatic double layer and make direct contact with the surface (Carniello et al., 2018). This effect might explain why WT spores showed stronger and more stable adhesion in our experiments. In a previous study, we used an optical tweezer system and found that WT spores could be moved along a surface, but not lifted off, suggesting that they were tethered by their appendages (Jonsmoen et al., 2023). The same study also showed that L-ENAs are especially important for anchoring spores to surfaces. In contrast, bald spores, which lack these appendages, do not have the same ability to attach directly to surfaces. This likely explains their weaker initial adhesion, especially to stainless steel (SS). However, when the ionic levels increase and the electrical double layer is compressed, the spore-surface might come in close range of the material surface which facilitates increased adhesion through other mechanisms than surface appendages (Carniello et al., 2018).

Acidic and caustic washes are commonly used in cleaning-in-place procedures to remove biological material residues from pipes and processing equipment. However, our finding suggests that spore adhesion of bald spores increases under both acidic and alkaline conditions compared to pure water, while the WT maintained its strong adhesion. This enhanced surface adhesion could potentially reduce the effectiveness of industrial cleaning processes, as spores adhered to surfaces tend to have an enhanced tolerance to alkali compared to spores that remain free in suspension (Nanasaki et al., 2010). In real-world processing environments, spores are often introduced though complex media, like milk which contains a variety of components such as salts, lipids and proteins. Our results show that the adhesive behavior of spores is strongly influenced dependent by the surrounding suspension. Specifically, spores suspended in surfactants (e.g., Tween-20), or in solubilized protein (e.g., BSA) show reduced adhesion, whereas higher ionic strength promotes stronger attachment. To effectively reduce spore adhesion during processing, more knowledge is needed about how spores behave in complex food matrices like milk. Understanding which factors limit or promote initial adhesion will be crucial for developing improved cleaning strategies and preventing contamination.

In conclusion, this study has specifically investigated the role of ENAs in spore adhesion to abiotic surfaces using WT spores and spores of a panel of isogenic mutants and vegetative cells. Our findings demonstrate that ENAs play a significant role in enhancing adhesion efficiency to materials such as PP and SS. Substrate surface hydrophobicity was found not to be a driving factor for spore adhesion, but rather properties such as surface roughness were highlighted. The importance of understanding spore adhesion is underscored by the fact that attached spores exhibit markedly increased resistance to various stressors. For instance, adhered spores can display increased alkali tolerance and up to a 400% increase in heat tolerance compared to non-adhered spores (Nanasaki et al., 2010; Simmonds et al., 2003). Such resilience poses challenges in both food processing environments and clinical settings, where spore contamination can lead to significant safety and hygiene concerns. Gaining a deeper understanding of the factors influencing spore adherence is critical for developing targeted strategies to mitigate spore contamination. This knowledge is essential for designing effective cleaning protocols, improving surface materials to reduce spore adhesion, and ultimately minimizing the risks associated with spore persistence in sensitive environments.

## Materials and methods

### Spore preparations

Spores of the *B. paranthracis* strain NVH 0075/95 (Lund & Granum, 1996) and its isogenic mutants with different appendage compositions were used in this study, in addition to an NVH 0075/95 Δ*exsY* mutant with a deficient exosporium. Comparing multiple strains enabled us to isolate the specific effects of the appendages, offering deeper insight into their role in adhesion to abiotic surfaces. The genotypes and corresponding phenotypes of all mutant strains are presented in Table 1.

The spores were prepared in a sporulation media containing 8 g/L Nutrient Broth (Oxoid™, Cat. No: CM0001, Thermo Scientific), 5 mM (NH4)2SO4, 1 mM MgCl2, 1 mM Ca(NO3)2, 1 µM FeSO4, 66 µM MnSO4, 12.5 µM ZnCl2, 2.5 µM CuCl2, 2.5 µM Na2MoO4 and 2.5 µM CoCl2, as adapted from van der Voort et al., 2010. The cultures were incubated at 37 °C in a conical flask with shaking at 200 rpm. Spores were harvested when ≥ 90% of the vegetative cells had sporulated, as determined by phase-contrast microscopy. This was followed by an initial 10-minute centrifugation at 3700 x g (Heraeus Megafuge 16R Centrifuge equipped with a TX-400 rotor, Thermo Scientific™), followed by three washes with sterile water (4 °C). The spores were stored at 4 °C in sterile water until use.

Fresh cultures of *B. paranthracis* were prepared by adding 1 mL of an LB overnight culture to 9 mL fresh LB media and incubated for 3 hours at 37 °C. The vegetative cells were harvested using the same procedure as for the spores, albeit with centrifugation at 2500 x g for 10 minutes.

### Adhesion experiments

Spore adhesion testing was performed using a modified version of the method described by Hamiot et al., 2023. Initially, the number of transfers was determined to be ten, based on the measurement of how many transfers were required to reach an OD of 0.5 for WT in a polystyrene cuvette. The spore concentration in a sterile water suspension was adjusted to 0.8 by measuring the optical density at 600 nm (OD600). A volume of 1 mL of the OD adjusted spore suspension was transferred to a PS cuvette (BR75901 5, Brand) using a Pasteur pipette (No. 612-1702, VWR), and the OD600 was measured again. Pasteur pipettes were used as spores exhibited lower adherence to the glass material of these pipettes, compared to plastic pipette tips, as determined by visual inspection (data not shown). This procedure was repeated nine additional times, with the spore suspension sequentially transferred to new cuvettes and OD600 recorded after each transfer.

The decrease in OD600 was used as an indirect measure of the number of spores that adhered to the vial surfaces. The following formula calculated the proportion of non-adherent spores remaining in the suspension: ODn/ODi, where ODi is the initial OD600 and ODn is the OD600 after n transfers. A similar procedure was also used to analyze spore adhesion to glass, SS, and PP tubes. Here, we used glass test tubes (14 mm X 130 mm, DWK Life Sciences), 316 stainless steel pipe fittings (No. 499-3467, RS PRO), and 1.5 mL microcentrifuge tubes of PP (REF 72.690.001, Sarstedt). Adhesion by vegetative cells was assessed using the same procedure as for spores for all materials tested. Lastly, the same setup was used to investigate how phosphate buffered saline (PBS), bovine serum albumin (1%, A6003, Sigma-Aldrich) and Tween-20 (0.05%, BP337-100, Fisher BioReagents) affected adhesion to SS by suspending the spores in the respective solutions, instead of sterile water.

### Assessing adhesion over time

The WT and isogenic mutant strains were prepared as described in the previous section on adhesion experiments by adjusting initial OD600 to 0.8. To observe the effect of adhesion over time the tubes were placed in a HulaMixer (15920D, Thermo Scientific) set to rotate at 40 rpm. They were allowed to rotate for either 1 minute, 10 minutes, 30 minutes or 1 hour. After each time point, the tubes were removed from the mixer and the OD600 was measured. Calculations were performed as described in the previous section.

### Microtiter experiments

To compare the role of material wettability on spore adhesion, we used microtiter plates to ensure uniform measurements. The plates used for this analysis included a PS Nunclon Delta-treated plate (168055, Thermo Scientific^TM^) representative of hydrophilic surface-treated polystyrene, and a hydrophobic COSTAR PS-untreated plate (Corning® Polystyrene Not Treated, Cat.no. CLS3370, Corning, Inc.). The OD600 of the spore suspension was adjusted to 0.8, and 200 μL was transferred to the first well. After pipetting the spore suspension into the first well, the plate was loaded into the Infinite M200 microplate reader (Tecan Trading AG). The OD600 of the sample was recorded whereafter the spore suspension was transferred 3-, 6-, and 9-times using Pasteur pipettes. Calculations were performed as described for the adhesion experiments section above.

### Microbial adhesion to hydrocarbon

The relative hydrophobicity of spores was estimated using the microbial adhesion to hydrocarbon (MATH) assay, as described in Jindal & Anand, 2018; Rosenberg, 2006. The spore suspension was diluted to an OD600 of approximately 1.6 in glass test tubes (14 mm X 130 mm, DWK Life Sciences) and a volume of 1.5 mL of hexadecane (99% purity, H6703, Sigma-Aldrich) was added to 2 mL of the spore suspension. The tubes were vigorously vortexed for 1 minute and left to settle at 30 °C before being vortexed for another 2 minutes. The phases were allowed to separate at room temperature for 20 minutes before the OD600 of the bottom aqueous phase was measured.

The percentage of relative hydrophobicity was calculated using the following formula:

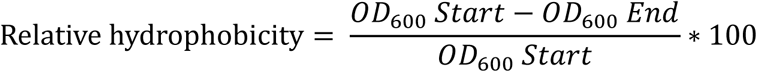

### Contact angle measurements

Contact angle measurements were performed using a Theta One tensiometer (Attension®, Biolin Scientific). A 4 µL drop of deionised water was generated above the material surface using a gauge 22 needle (C209-22) at a rate of 1 μL/s. A total of 150 image frames were acquired over 10 seconds immediately after the drop was deposited and the contact angle was determined.

### Surface roughness analyses

Surface topographies were characterized using AFM or stylus profilometry, depending on sample type. AFM scans (100 µm²) were acquired in tapping mode using a Park NX-Hivac (Park Systems Corp.) system with a PPP-NCHR (Park Systems Corp.) probe (tip radius <10 nm). SS surfaces were instead scanned using a Dektak XT stylus profilometer (Bruker) over 1 mm² areas with 10 µm lateral resolution, to better capture larger-scale surface features. Surface maps were processed in Gwyddion (v. 2.67) (Nečas & Klapetek, 2012) in order to minimize imaging artifacts, using sample-specific background correction procedures.

We quantified three key surface parameters, as an average of three measurements from different samples (two for glass): Root mean square (RMS) roughness, reflecting vertical height variation, surface ratio (SR = surface area / projected area), indicating topographical expansion, and autocorrelation length (ACL), describing the lateral scale over which surface features remain correlated. ACL was estimated using two complementary approaches. First, an orientation-independent ACL was derived from the 2D power spectral density function (PSDF). Second, directional ACLs were evaluated along horizontal and vertical axes using Gwyddion’s correlation length tool. In both cases the ACL was estimated based on the distance at which the autocorrelation function decays to 1/e of its maximum.

### Electron microscopy

Scanning electron microscopy was used to observe spore adhesion on SS, PP, PS and glass surfaces. The materials were cut into smaller pieces suitable for SEM analysis, then immersed in spore suspension (OD600 0.8) prepared in sterile water. After 1 hour of incubation, the spore suspension were fixed with a 4% formaldehyde solution, which was applied for 15 minutes. The materials were then rinsed three times with sterile water and left to dry overnight. Subsequently, the samples were coated with an approximately ∼30 nm layer of platinum using a sputter coater (Leica EM ACE200). Images were captured using a Zeiss EVO 50 scanning electron microscope operating in high vacuum mode with an accelerating voltage of 10kV and a current of 20 pA, and at a magnification of 3000x.

## Data analysis and statistics

Statistical analysis was performed using Prism (Prism 10.3, GraphPad). One-way ANOVA was applied to the data from the 10^th^ transfer in each experiment series, followed by Dunnett’s (Figure 2, 5, 7, 8 and 9) multiple comparisons test to check for differences relative to the WT strain, or Tukey’s (Figure 4) multiple comparisons test to assess differences between all strains. Statistical significance is indicated as follows; *p* > 0.05 (ns), *p* ≤ 0.05 (*), *p* ≤ 0.01 (**), *p* ≤ 0.001 (***) and *p* ≤ 0.0001 (****). All p-values are summarised and available in Supplementary Information under Statistical analysis results, tables S6-S11. Graphing and data visualization were performed using Origin 2024 (OriginLab).

## Author Contributions

Unni Lise A Jonsmoen: Conceptualization, investigation, methodology, writing – original draft, review, and editing. Jennie Ann Allred: investigation, methodology, writing – original draft, review, and editing. Dmitry Malyshev: methodology, writing – review and editing, Jonas Segervald: investigation, writing – review and editing, Magnus Andersson: conceptualization, writing – review and editing, funding acquisition, Marina Elisabeth Aspholm: conceptualization, funding acquisition, methodology, project administration, resources, writing – editing.

## Acknowledgements

The authors acknowledge the facilities and technical assistance of the NMBU Imaging Center. Ephrem Debebe Zegeye for supervision of ULJ. Special thanks to Yohannes Beyene Mekonnen, Kristin (Tina) O’Sullivan, Johan Jonsson and Daniel P. G. Nilsson for technical assistance. This work was supported by the Norwegian Research Council (33529) and the Norwegian University of Life Sciences (NMBUs) research fund to M.E.A., and the Swedish Research Council (2023-04085) to M.A.

## Conflict of interest statement

The authors declare no conflicts of interest.

